# Communication of perceptual predictions from the hippocampus to the deep layers of the parahippocampal cortex

**DOI:** 10.1101/2024.03.28.587186

**Authors:** Oliver Warrington, Nadine N. Graedel, Martina F. Callaghan, Peter Kok

## Abstract

Current evidence points to the hippocampus as an essential region coordinating learning and exploiting predictive relationships in the service of perception. However, it remains unclear whether the hippocampus drives the communication of predictions to the sensory cortex or acts as a recipient of predictions from elsewhere. Here, we collected sub-millimetre 7T fMRI data to investigate neural signals in the medial temporal lobe (MTL). We used layer-specific fMRI to infer the direction of communication between the hippocampus and cortex. Specifically, superficial layers of the MTL cortex project to the hippocampus, while deep MTL layers receive feedback projections. Participants performed a task in which auditory cues predicted abstract shapes. Crucially, we omitted the expected shape on 25% of trials, thus isolating the prediction signal from bottom-up input and allowing us to ask: In which direction are predictions communicated between the hippocampus and neocortex? Neural patterns in CA23, pre/parasubiculum and the parahippocampal cortex (PHC) reflected shape-specific predictions. Layer-specific informational connectivity analyses revealed that communication between CA23 and PHC was specific to the deep layers of PHC. These findings are in line with the hippocampus generating predictions through pattern completion in CA23 and feeding these predictions back to the neocortex.

## Introduction

The brain must use prior knowledge to infer the cause of incoming sensory information. Prior knowledge informs predictions of future sensations, relying on our ability to extract statistical regularities from the environment, including rapidly learnt associations between arbitrary stimuli. While the effects of predictions on sensory processing in the early sensory cortex are evident [3], the mechanisms underlying the generation and communications of these predictions remain unclear. The hippocampus has recently been suggested as an essential region coordinating learning and exploitation of predictive relationships for perceptual inference [3,4]. Representations of predicted stimuli have been found in the hippocampus [5–7], and informational connectivity between the hippocampus and visual cortex increases after an association has been learnt [8,9]. However, with typical fMRI analyses, these studies could not determine whether the hippocampus was responsible for passing these predictions to the cortex. One possible mechanism to link these findings is that the hippocampus generates and communicates predictions to the sensory cortex via mechanisms used in episodic memory, such as pattern completion and cortical reinstatement [6,10–13]. Indeed, predictions could exploit the reversal of information flow between the sensory cortex and the medial temporal lobe that has been found for internally-generated information in memory [14–19] and mental imagery [20].

Here, we used 7T layer-specific fMRI to determine the direction of communication between the hippocampus and neocortex when a prediction signal was isolated from bottom-up sensory input. The key to inferring the direction was separating the layers of the medial temporal lobe (MTL) cortex, i.e. entorhinal (ERC), parahippocampal (PHC) and perirhinal (PRC) cortices. ERC layers II and III project to the hippocampus, while layer V receives feedback projections [21]. Analogously, superficial layers of PHC and PRC project to ERC and onward to the hippocampus, whereas deep layers of PHC and PRC receive feedback from the hippocampus via ERC. In short, feedback from the hippocampus to the neocortex would be expected to be reflected specifically in the deep layers of the MTL cortex. Participants performed a task in which an auditory cue predicted which one of two possible shapes would be presented. The predicted shape was shown in 75% of the trials, but crucially, we omitted the shape in the remaining 25%, thus isolating the prediction signal from the bottom-up input.

## Results

### Representations of predicted but omitted shapes in the hippocampus

With the improved spatial resolution of 7T fMRI, we could test our hypotheses about the involvement of the hippocampus in prediction signalling at different levels of anatomical subdivisions, from the whole hippocampus to longitudinal and subfield-specific splits. At the subfield level, we hypothesised that information about predicted shapes would be present in CA3 after the auditory cue had triggered pattern-completion of the full event [5–7]. We also hypothesised that this representation would be reflected in the subiculum as it travelled from the hippocampus to the cortex [7,8]. Previous research has found a change in the complexity of representations along the long axis of the hippocampus [24], and that the posterior hippocampus is involved in signalling perceptual predictions and prediction errors based on simple associations [8,9,25]. Therefore, we predicted that the posterior hippocampus and its subfields would contain information about the expected shape.

To test for hippocampus involvement in prediction signalling, we used a multi-voxel correlation analysis [26] to investigate the pattern similarity of shape-specific BOLD responses between presented and predicted-but-omitted shapes (Figure 1 B). To determine shape-specific activity patterns, activity evoked by shape B was subtracted from activity evoked by shape A in each voxel. We then calculated the Pearson’s R correlation between shape-specific activity patterns across voxels from the localiser run, in which shapes were presented but not predicted, and shape-specific activity patterns from the four prediction runs, where shapes were predicted but omitted. Before running statistical tests, we converted Pearson’s R values to Fisher’s Z.

**Figure 1:**
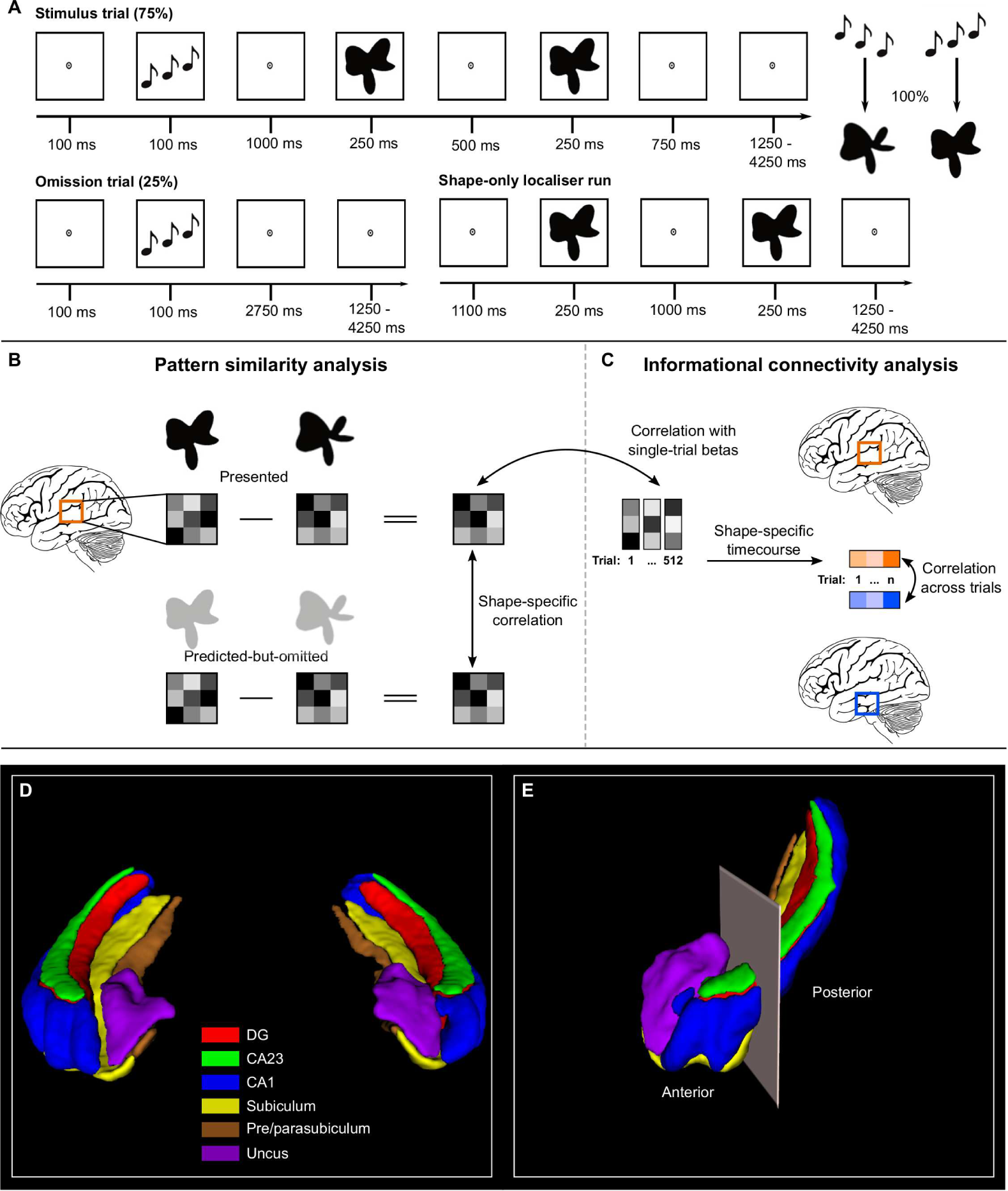
Overview of experimental paradigm and analysis. A) Prediction runs include stimulus and omission trials. In stimulus trials, the auditory cue preceded the presentation of two consecutive shape stimuli. On each trial, the second shape was identical to the first or slightly warped along an orthogonal dimension. The participants’ task was to report whether the two shapes were the same or different. During prediction runs, the auditory cue (ascending vs descending tones) predicted whether the first shape on that trial would be A or B. The cue was valid in 75% of trials (stimulus trials), whereas the expected shape was omitted in the other 25% of trials (omission trials). On omission trials, participants had no task and were told to wait for the subsequent trial to start. We had a single shape-only localiser run in which participants performed the same shape-discrimination task but had no auditory cue. The separate localiser run allowed us to contrast representations of the same shape when presented or predicted but omitted. B) To perform the pattern similarity analysis, the activity patterns of shape B were subtracted from shape A separately for presented shapes and predicted-but-omitted shapes. We then calculated the Pearson’s R correlation between presented and predicted but omitted shapes for each region of interest. C) To perform the informational connectivity analysis, the shape-specific activity pattern from the localiser was correlated with single-trial betas from the omission trials for each region to generate a shape-specific timecourse of information. We then correlated these timecourses between regions to determine their shared fluctuations in shape-information. D) 3D render of a representative hippocampal subfield segmentation of both hemispheres viewed from the anterior perspective. E) 3D render of the anterior-posterior boundary viewed from the anterolateral perspective. The boundary was defined as the last coronal slice in which the uncus was visible. Segmentations were generated with the ASHS toolbox [22] trained on an atlas of manual 7T segmentations [23].

At the gross anatomical level, we found that the hippocampus as a whole and the posterior hippocampus showed a trend for a negative correlation between presented and predicted-but-omitted shapes (whole: Z = −0.069, t_29_ = −2.00, p = .055; posterior: Z = −0.086, t_29_ = −1.95, p = .061)(Figure 2), while there was no evidence of anterior hippocampus involvement (Z = −0.049, t_29_ = −1.32, p = .199). The negative correlation was surprising as we expected shape predictions to evoke similar activity patterns as presented shapes. Due to the subtraction logic of our analysis, a negative correlation could reflect either a prediction evoking a negative image of the pattern evoked by the presentation of the same shape or a representation of the non-prediction shape (i.e. shape A cued but shape B represented). To distinguish between these explanations, we separately calculated within-shape (e.g. between shape A predicted and shape A presented) and between-shape correlations (e.g. between shape A predicted and shape B presented). For both the hippocampus and the posterior hippocampus, only within-shape correlations were significantly different from 0 (hippocampus within: Z = −0.048, t_29_ = −2.50, p = .018; between: Z = −0.010, t_29_ = −0.44, p = .662; posterior hippocampus within: Z = −0.066, t_29_ = −2.84, p = .008; between: Z = −0.018, t_29_ = −0.73, p = .469) and within-shape correlations were significantly more negative than between-shape correlations (hippocampus difference: Z = −0.038, t_29_ = −2.33, p = .027; posterior hippocampus difference: Z = −0.048, t_29_ = −2.15, p = .04)(Figure 2). These results show that activity patterns evoked by predicted but omitted shapes are opposite in sign to those evoked by presented shapes. These signals could reflect either inhibitory prediction signals [4] or negative prediction errors [1,27].

**Figure 2:**
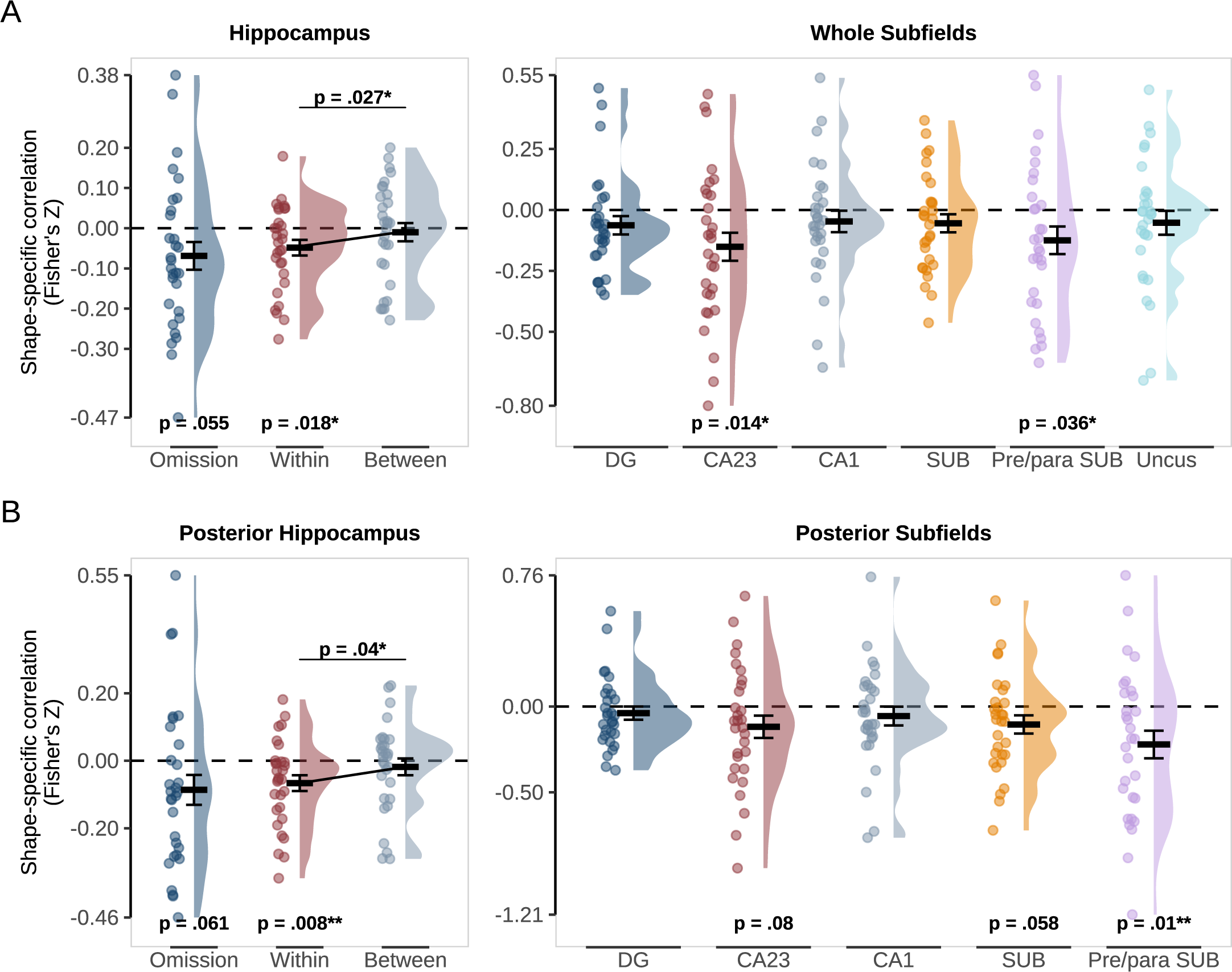
Pattern similarity between predicted but omitted shapes and presented shapes in the hippocampus. A) Whole Hippocampus (left) and subfields (right). B) Posterior Hippocampus (left) and subfields (right). Omission represents similarity in shape-specific (shape B - shape A) activity patterns in the localiser and omission trials with p values for a two-sided, one-sample t-test against 0. Within and Between reflect pattern similarity within the same shape (e.g. shape A presented - shape A omitted) and between shapes (e.g. shape A presented - shape B omitted) with p values for a two-sided, paired t-test. Crossbars and error bars represent the mean and within-subject standard error of the mean, respectively. Individual subject values are plotted in points alongside the probability density estimate.

Given these results, we restricted our subfield-specific investigations to whole and posterior subfields. Sub-fields were segmented using ASHS [22] with an atlas based on manual segmentation of 7T anatomical scans [23]. This revealed prediction information in CA23 (whole: Z = −0.139, t_29_ = −2.63, p = .014; posterior Z = −0.106, t_29_ = −1.81, p = .08) and the pre/parasubiculum (whole: Z = −0.116, t_29_ = −2.22, p = .034; posterior: Z = −0.189, t_29_ = −2.78, p = .009). The subiculum was at trend level only in the posterior hippocampus (whole: Z = −0.548, t_29_ = −1.48, p = .15; posterior: Z = −0.105, t_29_ = −1.98, p = .058), while no other subfields contained shape-specific representations of predicted-but-omitted shapes (whole: all p > .1; posterior: all p > .3). It is worth noting that the pre/parasubiculum was not distinguished from the subiculum proper in previous 3T fMRI work investigating prediction signals in the hippocampus [5,7,8]. Indeed, segmenting the hippocampus with the 3T atlas used previously [28,29] revealed strong prediction signals in the posterior subiculum in the current dataset (Z = −0.19, t_29_ = −3.60, p = .001), consistent with previous work [7,8]. This finding raises the possibility that the pre/parasubiculum was the driving force behind previous prediction signals in the subiculum. We report the 3T atlas results in the supplemental information (Supplemental table 1).

### Parahippocampal Cortex contains predicted but omitted shape information

If predictions are communicated between the hippocampus and neocortex, we hypothesised that prediction representations would be found in the cortical regions of the MTL. To test this hypothesis, we extended the pattern similarity analysis to ERC, PRC and PHC. We hypothesised that we would find representations of predicted shapes in ERC, a major target of hippocampus output, and that these representations would be specific to the deep layers [21]. As the visual cortex is presumably the ultimate target of a shape prediction signal, we also hypothesised that PRC, PHC, or both would be involved as the signal passed down the cortical hierarchy to the sensory cortex.

We found evidence for activity patterns reflecting predicted-but-omitted shapes in PHC (Z = −0.073, t_29_ = −2.18, p = .037) but not in ERC or PRC (both p > .3). In PHC, the prediction signal was driven by a statistically significant representation in the deep and middle but not superficial layers (deep: Z = −0.094, t_29_ = −2.72, p = .011; middle: Z = −0.075, t_29_ = −2.04, p = .05; superficial: Z = −0.083, t_29_ = −1.73, p = .095)(Figure 3 A). A repeated-measures one-way ANOVA revealed no significant difference between layers (F_(2,29)_ = 0.17, p = .844), meaning we cannot draw conclusions about the layer-specificity of this effect. However, note that an effect in the deep layers cannot be explained by the draining vein effect common to layer-specific BOLD fMRI [30], suggesting at the least a genuine involvement of PHC deep layers in prediction signalling. Consistent with the lack of an effect in ERC as a whole, we did not find evidence for a prediction signal in ERC in any layer (all p > .3).

**Figure 3:**
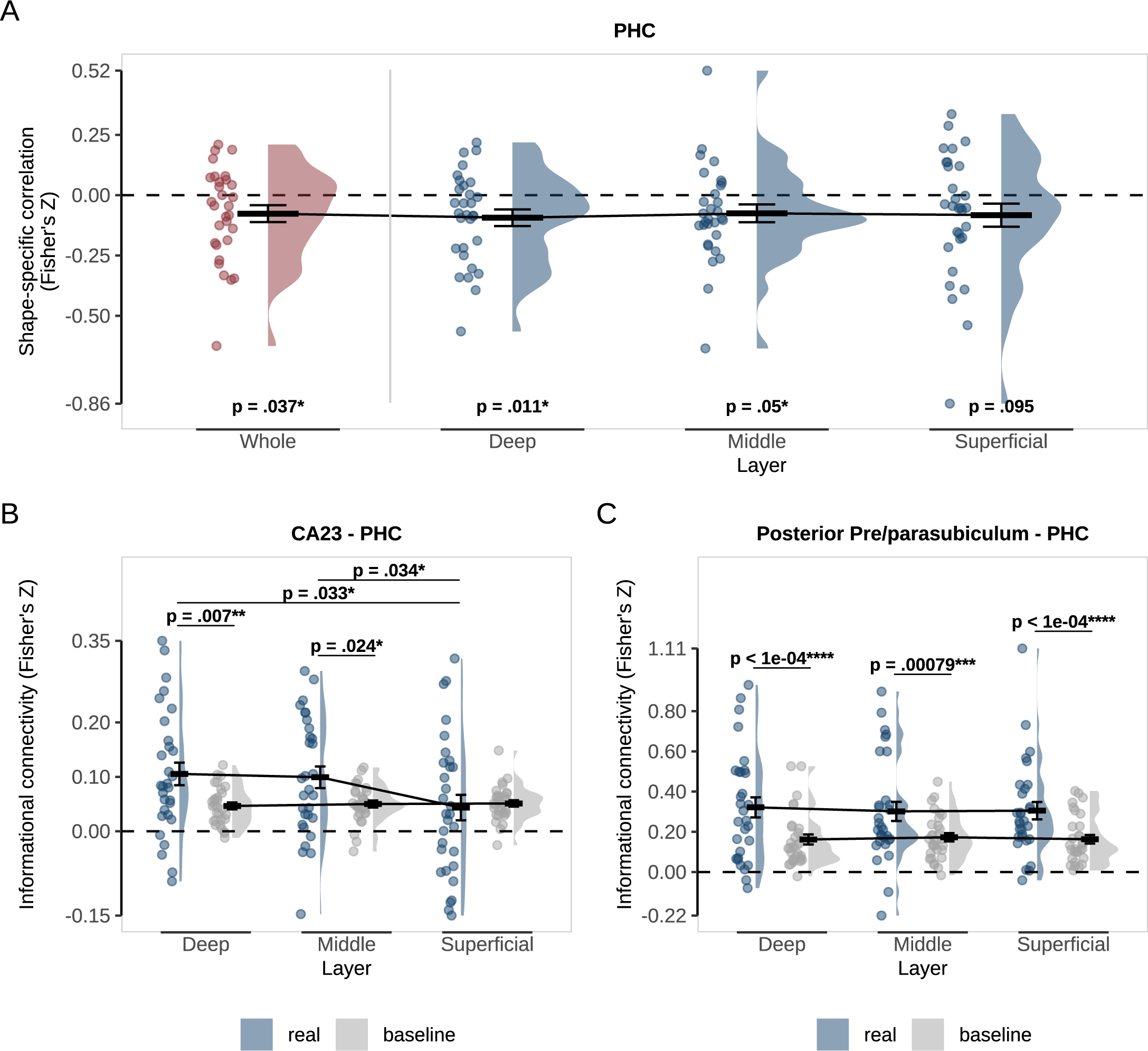
Representations and connectivity of PHC during omission trials. A) Pattern-similarity analysis in PHC as a whole (red) and specific to the deep, middle and superficial layers (blue). Pattern-sim-ilarity reflects the correlation between shape-specific (shape B - shape A) activity patterns in the localiser and omission trials with p values for a two-sided, one-sample t-test against 0. B) Informational connectivity of CA23 and C) pre/parasubiculum with PHC layers. Real connectivity (blue) is the observed correlation between regions on omission trials. Baseline connectivity (grey) was calculated by randomly shuffling the shape labels across 100 permutations. p values represent post-hoc paired t-tests investigating the differences between real and baseline and across layers. For all figures, crossbars and error bars represent the mean and within-subject standard error of the mean, respectively. Individual subject values are plotted in points alongside the probability density estimate.

### Informational connectivity between the hippocampus and cortex during predictions

Having established that prediction signals are evident in the hippocampus and PHC, we tested our main research question - What is the direction of communication between the hippocampus and cortex during prediction signalling? To answer this question, we calculated informational connectivity between the layers of PHC and the hippocampal subfields that contained significant prediction signals: CA23 and pre/para-subiculum. Informational connectivity is a multivariate version of functional connectivity that determines the correlation between pattern-similarity measures across time between two regions [8,31,32].

To calculate the pattern-similarity time course for each ROI, we correlated the activity pattern resulting from the shape A - shape B contrast in the localiser with single-trial activity patterns in the omission trials. We also calculated scrambled pattern-similarity time courses by shuffling shape labels across 100 permutations to compare “baseline” and “real” connectivity. To obtain subfield specificity, despite their spatial contiguity, we calculated the partial correlation between a subfield of interest and PHC layers by regressing out all other subfields (see Methods for details). Connectivity values for each subject were entered into a two-way repeated-measures ANOVA to test for an interaction between connectivity (real, baseline) and layer (deep, middle, superficial). Like univariate functional connectivity, informational connectivity alone cannot determine the direction of communication. However, by combining the connectivity measure with layer-specific fMRI, we can use the known anatomical connections of MTL to infer directionality [33].

Strikingly, PHC showed differential connectivity across layers with CA23 (F_(2,58)_ = 5.09, p = .009)(Figure 3 B). Follow-up t-tests showed connectivity greater than baseline in deep and middle but not superficial layers (deep: t_29_ = 2.89, p = .0072; middle: t_29_ = 2.38, p = .0239; superficial: t_29_ = −0.33, p = .7455) with both deep and middle being significantly above superficial (deep vs superficial: t_29_ = 2.24, p = .0328; middle vs superficial: t_29_ = 2.22, p = .0341; deep vs middle: t_29_ = 0.39, p = .6982). Pre/parasubiculum, on the other hand, revealed a main effect of connectivity with PHC (whole: F_(2,58)_ = 19.45, p < .001; posterior: F_(2,58)_ = 25.62, p < .001)(Figure 3 C), but no significant difference across layers (both p > .6). Given our original hypothesis that ERC would be involved, we also probed its connectivity with hippocampal subfields. There was a main effect of connectivity between ERC and Pre/parasubiculum (whole subfield: F_(2,58)_ = 8.29, p = .007; posterior subfield: F_(2,58)_ = 10.98, p = .002), but not CA23 (F_(2,58)_ = 3.83, p = .060). There was no interaction between connectivity and layer for either subfield (both p > .8).

Finally, in an exploratory analysis, we investigated connectivity to the shape-selective Lateral Occipital Cortex (LOC) [34], a region whose representations have previously been shown to co-vary with hippocampal predictions across subjects [7]. This analysis revealed layer-specific connectivity between LOC and posterior CA23 (posterior subfield: F_(2,58)_ = 3.16; p = .049; whole: F_(2,58)_ = 1.19, p = .313), but not pre/parasubiculum (all p > 0.9). Follow-up t-tests showed that connectivity was not significantly different between real and baseline in any layer, though there was a trend in the deep layers (deep: t_29_ = 1.86, p = .072; middle: t_29_ = 1.10, p = . 278; superficial: t_29_ = −0.03, p = .974). Deep and middle layers showed significantly stronger connectivity with posterior CA23 than superficial layers (deep vs superficial: t_29_ = 2.26, p = .031; middle vs superficial: t_29_ = 3.00, p = .005; deep vs middle: t_29_ = 1.07, p = .296)(Figure 4) This exploratory analysis requires further replication and corroboration.

**Figure 4:**
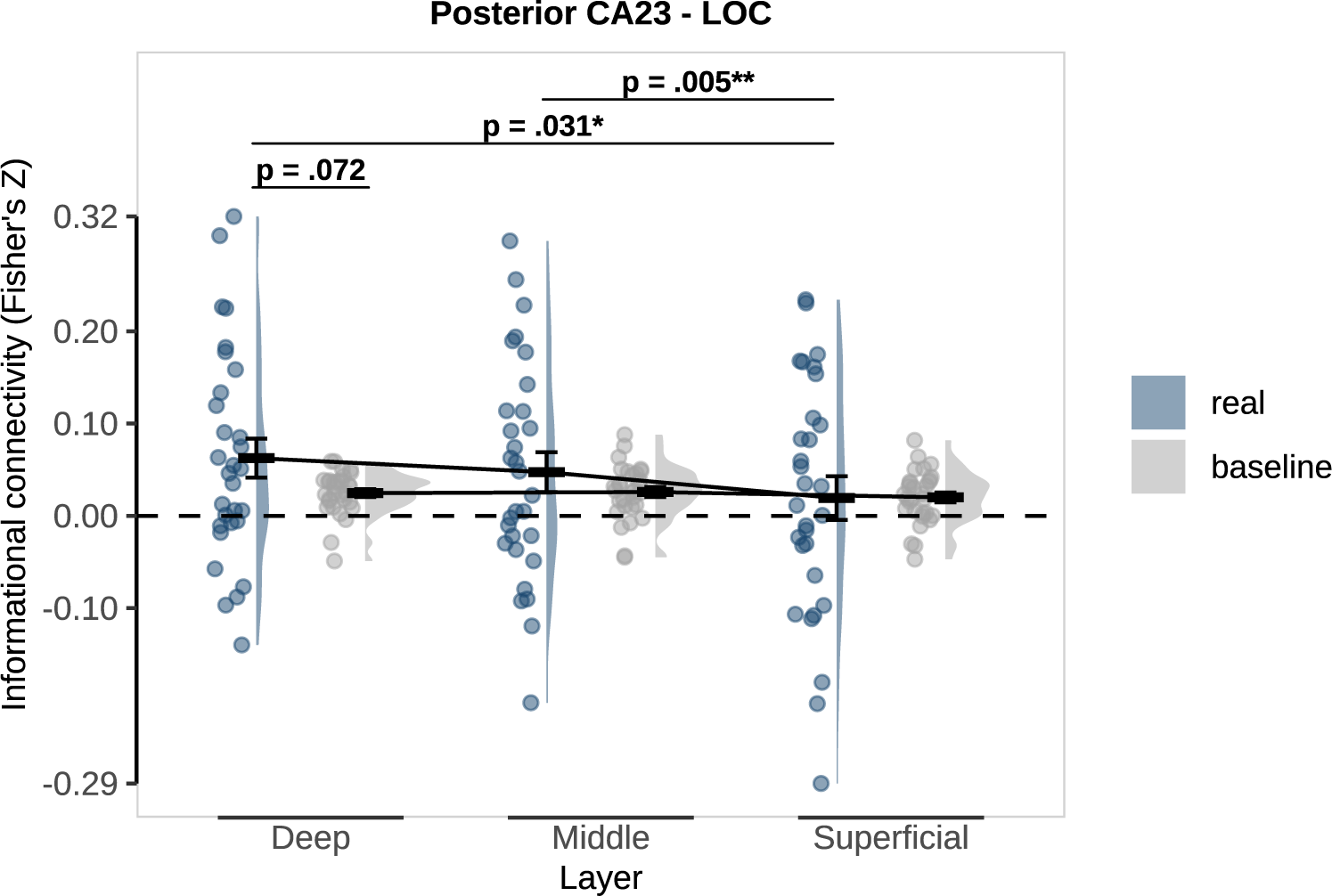
Layer-specific informational connectivity between posterior CA23 and LOC during omission trials. Real connectivity (blue) is the observed correlation between regions on omission trials. Baseline connectivity (grey) was calculated by randomly shuffling the shape labels across 100 permutations. p values represent post-hoc paired t-tests investigating the differences between real and baseline and across layers. Crossbars and error bars represent the mean and within-subject standard error of the mean, respectively. Individual subject values are plotted in points alongside the probability density estimate.

## Discussion

In summary, we found that after learning an arbitrary, multi-modal association, omission of the predicted shape led to the representation of shape-specific prediction signals in CA23, pre/parasubiculum and PHC. The representations of predicted shapes were negatively correlated with the representation of the same shapes presented in the absence of a prediction. We used layer-specific fMRI combined with informational connectivity analyses to determine the direction of communication between the hippocampus and neocortex during prediction signalling. We found evidence suggesting predictions are communicated from CA23 to the deep layers of PHC and possibly back to the shape-selective visual cortex.

Recently, Barron et al.[4] have incorporated the hippocampus into the predictive coding framework, in which feedback projections descend the cortical hierarchy to suppress the components of sensory input that can be correctly predicted [27,35,36]. As the hippocampus sits at the top of the sensory hierarchy [37], it acts as a hub region positioned high within a hierarchical generative model. From this position, it can generate predictions that “explain away” predictable input distributed across the sensory cortices.

According to this proposal[4] and previous suggestions [6,7], perceptual predictions rely on the same neural machinery underlying episodic memory. In particular, pattern completion is a vital function of the hippocampus that enables the retrieval of associated items from memory based on partial information [10,11]. Pattern completion has been attributed to CA3 [10,12,38,39] and previous studies have found evidence in line with CA3 representing predicted stimuli [6,7] but were unable to separate CA3 from DG, a subfield key for the opposing process of pattern separation [40,41]. In this study, we could define a separate CA23 ROI due to the increased spatial resolution possible with 7T fMRI, and we found evidence of stimulus-specific prediction signals in CA23 but not DG.

The principal aim of the current study was to determine the direction of predictive communication between the hippocampus and neocortex. To this end, we investigated the layers of MTL cortical regions, because the separation of feedforward and feedback information processing into largely different layers [37] allows inferences of directionality to be made [33]. Our preregistered hypothesis centred on ERC, as it is a major target of hippocampus output [21], and previous layer-specific fMRI studies were able to dissociate memory encoding and retrieval in the superficial and deep layers of ERC, respectively [42,43]. Recently, results from our group have also shown increased functional connectivity between the posterior subiculum and ERC once learning of a predictive association was complete [8]. However, we found no evidence of predicted shape representations in ERC. Likewise, there was no layer-specific informational connectivity between ERC and hippocampal subfields (Supplemental figure 4). These null results may suggest that ERC is not involved in signalling predictions, and perhaps other cortical regions communicate directly with the hippocampus, by-passing ERC [44,45]. However, it is important to note that ERC (and PRC) suffered greater drop-out and have lower temporal signal-to-noise than hippocampus and PHC (Supplemental figure 1), meaning our null results may be due to insufficient power.

In line with this, we did find information for the predicted-but-omitted shape in PHC, with layer-specific analyses showing this information was represented most robustly in the deep layers (Figure 3 A). We also found evidence of informational connectivity between CA23 and the deep layers of PHC during omission trials (Figure 3 B). Taken together, we interpret these results as information retrieved during pattern completion by CA23 sent from the hippocampus to the cortex. As far as we know currently, there is no direct connection between CA23 and PHC, and this functional relationship is likely mediated by subfields responsible for sending information to the cortex, such as CA1 and the subicular complex.

The pre/parasubiculum also showed a shape-specific prediction signal (Figure 2) and informational connectivity with PHC (Figure 3 C). However, the connectivity was not specific to the deep layers and was present across all layers of PHC instead. The involvement of all layers suggests ongoing bidirectional communication between pre/parasubiculum and PHC despite the omission of the bottom-up input. While the subiculum proper is the primary output subfield of the hippocampus [21,46,47], the pre/parasubiculum plays a dual role, both sending information out of the hippocampus but also acting as the primary target for visuospatial information sent to the hippocampus [48–53]. Therefore, it is unclear exactly what purpose the communication between pre/parasubiculum and PHC serves during omission trials. However, a speculative explanation may reflect an extension of the big-loop [11,32,54,55] in which hippocampal output about the predicted stimulus is returned as new input. Future studies will be needed to tease apart the precise nature of the bidirectional connectivity of pre/parasubiculum and PHC during predictions.

Although we hypothesised the involvement of pattern completion and cortical reinstatement in prediction signalling, we did not expect the regions involved to show a negative correlation between predicted and presented shape representations. This negative relationship was present in CA23, pre/parasubiculum and the PHC. It is unclear why predictions evoke a negative image of the pattern evoked during presentation. However, we offer a speculative explanation in the following.

When integrating the hippocampus into the predictive processing framework, Barron et al.[4] noted that information retrieved by pattern completion should have opposing influences on the cortex in memory and prediction. While the hippocampus should facilitate the cortex to reinstate episodic memories, it should instead suppress and explain away ascending cortical inputs during perception by sending inhibitory prediction signals. Therefore, the negative prediction representation we observe in this study may reflect an inhibitory prediction signal generated in the hippocampus and sent to the PHC to suppress the ascending shape information. Since we omit the predicted shape, this inhibitory prediction is not integrated with bottom-up input and remains detectable.

A negative prediction error is another possible explanation for the negative correlation [1]. In predictive-coding frameworks, two kinds of prediction errors are expected to exist. A positive prediction error would signal that an unexpected stimulus has appeared. In contrast, a negative prediction error occurs when a stimulus unexpectedly disappears or the expected stimulus does not appear, as in this study. Therefore, unpredicted shapes (in the localiser) and predicted-but-omitted shapes may activate separate populations of prediction error neurons, yielding a negative correlation. However, positive and negative prediction error neurons would be expected to be located in close proximity, and it is not clear that we would be able to pick up such separate populations with fMRI voxels.

As the BOLD signal can reflect excitatory and inhibitory signals, it is impossible to investigate these suggestions in the current dataset. However, new imaging methods that allow simultaneous measurement of brain activity and changes in glutamate and GABA concentrations are becoming available [56–58]. Koolschijn et al.[59] have already demonstrated the potential of concurrent fMRI and functional magnetic resonance spectroscopy (fMRS) by showing that hippocampal activity predicts an increase in the ratio of glutamate and GABA in the visual cortex during the recall of a visual cue from a paired associate. A future study using concurrent fMRI-fMRS could determine whether the hippocampus is associated with increased inhibition during prediction signalling or whether predictions rely on a similar disinhibitory mechanism as memory recall.

In conclusion, the current findings provide evidence that the hippocampus plays an important role in exploiting the regularities of our environment during perceptual prediction signalling and highlight the importance of further investigation into the role of the MTL cortex, particularly PHC, in perception. These findings add weight to suggestions that cognitive functions involving the internal generation of information, such as memory, planning and perception, may rely on similar mechanisms and, ultimately, the hippocampus.

## Methods

This study was preregistered on AsPredicted.org (https://aspredicted.org/F7J_GKJ).

### Participants

Thirty-five healthy human volunteers with normal vision and hearing participated in the study. Participants were excluded if they scored below 60% correct on trials with an in-time response or failed to respond to 50% or more of the trials. These measures were calculated across all fMRI runs of the session. Participants were also excluded if they failed to meet strict head motion criteria of maximum 10 movements of 1 mm or greater between successive functional volumes. Based on the above criteria, 5 participants were excluded from the final data set. The study was approved by the University College London ethics committee (approval number 8231_001). Participants were paid £8 for the behavioural session and £10 per hour for the fMRI session.

### Stimuli

Visual and auditory stimuli were generated using Matlab 2019a (Mathworks, Natick, MA, USA) and Psychophysics Toolbox 3.0.16 [60]. In the MRI scanner, visual stimuli were displayed on a rear projection screen using an Epson EB-L1100U projector (1920 × 1200 resolution, 60 Hz refresh rate) against a grey background. Participants viewed the visual display through a mirror that was mounted on the head coil with a viewing distance of 91 cm. An Ear-Tone Etymotic stereo sound system was used to play audio during scanning. The visual stimuli consisted of complex shapes defined with radial frequency components (RFCs). The contours of the stimuli were defined by seven RFCs, and a one-dimensional shape space was created by varying the amplitude of three out of the seven RFCs. Specifically, the amplitudes of the 1.11 Hz, 1.54 Hz and 4.94 Hz components increased together, ranging from 0 to 36 (first two components), and from 15.58 to 33.58 (third component). Note that we chose to vary three RFCs simultaneously, rather than one, to increase the perceptual (and neural) discriminability of the shapes. Five shapes were selected from this continuum such that they represented a perceptually symmetrical sample of this shape space, see [7] for details. From these 5 shapes, we selected shape 2 and shape 4, referred to as shape A and B in this study. A fourth RFC (the 3.18 Hz component) was used to create slightly warped versions of the five shapes, to enable the same/different shape discrimination cover task (see below). Black shapes subtending 4.5° were presented centered on fixation.

### Task

Participants performed five runs of the shape-discrimination task split into four prediction runs followed by a single shape-only localiser run (Figure 1 A). In the prediction runs, each trial began with a fixation bullseye presented for 100ms, followed by an auditory cue (ascending or descending tones) for another 100ms. After a 1000ms delay, two consecutive shape stimuli were presented for 250 ms each, separated by a 500 ms blank screen. The auditory cue predicted whether the identity of the first presented shape would be shape A or B. The cue was valid on 75% of trials (stimulus trials), whereas in the other 25% of trials, the predicted shape would be omitted entirely (omission trials). These omission trials isolate the top-down expectation signal from bottom-up stimulus input.

On each trial, the second shape was identical to the first or slightly warped. This warp was achieved by modulating the amplitude of the 3.18 Hz RFC component defining the shape. This modulation could be either positive or negative (counterbalanced over conditions), and the participant’s task was to indicate whether the two shapes on a given trial were the same or different using an MR-compatible button box. After the response interval ended (750 ms after the disappearance of the second shape), the fixation bullseye was replaced by a single dot, signalling the end of the trial while still requiring participants to fixate. This task was designed to avoid a direct relationship between the perceptual prediction and the task response. Furthermore, by modulating one of the RFCs not used to define our one-dimensional shape space, we ensured that the shape change on which the task was performed was orthogonal to the changes that defined the shape space and, thus, orthogonal to the prediction cues. An adaptive staircase procedure determined the modulation size [61] and was updated after each trial to make the task challenging (∼75% correct). The staircases were initialised in the behavioural session and continued throughout the experiment. However, if participants could not perform the task successfully in the scanner, the staircase was reset to maintain the task at a difficulty level that would maintain attention.

### Procedure

Participants attended two separate sessions within five days of each other. In the first session, participants were given instructions on the shape-discrimination task used throughout both sessions. The instructions were followed by 16 practice trials and one block of 128 trials, both shape-only trials with no auditory cues. To pass the threshold for inclusion, participants had to score > 60% correct on a single shape-only block and could repeat until their performance passed the threshold. Participants who succeeded were invited to the fMRI session and instructed on the prediction runs. The final part of the behavioural session was 32 practice trials to acclimatise to hearing the auditory cues during the task. The cues in the practice trials were 100% predictive of the identity of the first shape on that trial. For the first 15 participants, a coding error resulted in only shape A being presented in the practice runs. Control analyses did not reveal a significant difference in fMRI results between these participants and the subsequent 15 participants for whom this error had been corrected.

The second session was the fMRI scan. Participants performed a short reminder run of the shape-only trials during the whole-brain echo-planar imaging (EPI) acquisition (64 trials, 4 min). Participants were trained on the cue-shape associations in the scanner during practice runs that took place immediately before the prediction runs. That is, before the first prediction run, participants performed a practice run consisting of 64 trials, in which the auditory cue was 100% predictive of the identity of the first shape on that trial (e.g., ascending tones always followed by shape A and descending tones followed by shape B). Halfway through the experiment, the contingencies between the auditory cues and the shapes were flipped (e.g., ascending tones now followed by shape B and descending tones by shape A), and participants performed another practice run (64 trials, 4 min) to learn the new contingencies. The order of the cue-shape mappings was counterbalanced across participants. This procedure equated the frequencies of all tones and shapes and their transitions and ensured that stimulus differences could not explain any differences between valid and invalid trials. The two practice runs took place while anatomical scans (see below) were acquired to use scanner time fully. Finally, participants completed one shape-only run (128 trials) with no auditory cues.

### MRI acquisition

MRI images were acquired on a Siemens 7T Magnetom Terra (Siemens Healthcare GmbH, Erlangen, Germany) at the Wellcome Centre for Human Neuroimaging (University College London). Partial-brain functional images were collected with a T2*-weighted gradient-echo 3D EPI protocol (volume acquisition time = 3432 ms, TR = 39.00 ms, TE = 19.50 ms, water-selective excitation (1-2-1 binomial pulse, with 500 ms inter-pulse interval) with flip angle 13 degrees, voxel size 0.92 x 0.92 x 0.92 mm^3^, field of view 192 x 192 x 80.96 mm^3^, acceleration factor 4 in-plane and acceleration factor 2 in the second phase-encoded direction, in-plane segmentation factor 2, in-plane partial Fourier 6/8, echo spacing 1.20 ms, transverse slab with posterior-anterior phase-encoding direction). Four EPI volumes with reversed phase-encoding polarity was acquired immediately prior to the acquisition of the fMRI data to facilitate correction of susceptibility-induced distortions [62]. Anatomical images were acquired using an MP2RAGE sequence (TR = 5000 ms, TE = 2.54 ms, TI = 900 ms and 2750 ms, flip angles of 5° and 3°, voxel size 0.65 mm isotropic, field of view 220 x 220 x 156 mm^3^, GRAPPA acceleration factor 3) and a high-resolution T2-weighted sequence (TR = 3500 ms, TE = 229 ms, voxel size 0.52 x 0.52 x 0.50 mm^3^, field of view 168.8 x 200 x 56 mm^3^, GRAPPA acceleration factor 2). A whole-brain MT-weighted EPI image with distortions matched to the functional acquisition was acquired to aid with the coregistration of the anatomy to the functional data [62]. A second volume was acquired with reversed phase-encoding polarity.

### Anatomical preprocessing

#### Cortical surface reconstruction and coregistration

Accurate cortical surface reconstruction is required to define the cortical layers. In order to maximise the quality of the automated pipeline and thus minimise the manual edits required, several preprocessing steps were applied to the MP2RAGE data before submission to the Computational Anatomy Toolbox (CAT) [63] and FreeSurfer (https://surfer.nmr.mgh.harvard.edu). First, the gradient-echo image acquired at the later inversion time (INV2) volume was bias-corrected and segmented using the unified segmentation method available in Statistical Parametric Mapping (SPM12, http://www.fil.ion.ucl.ac.uk/spm, Wellcome Centre for Human Neuroimaging, London, UK). The CSF, bone, non-brain tissue and background tissue classes were combined with a threshold of > 0.5 and then inverted to create a brainmask. This brain mask provided the most accurate removal of dura in the visual cortex and around ERC. However, manual edits were still required around ERC as no automated method to date seems able to handle this strip of the dura consistently. The combined uniform (UNI) volume was denoised with the mp2rage toolbox for SPM12 (https://github.com/benoitberanger/ mp2rage) with a threshold of 6. The denoised UNI image was then skullstripped using the INV2-derived brainmask. The pial and white matter surfaces were reconstructed with both CAT and FreeSurfer. The pial surface generated by CAT in ERC was more consistently accurate than FreeSurfer; however, in the visual cortex, FreeSurfer outperformed CAT. Therefore, we generated layers using both programmes, CAT for MTL ROIs and FreeSurfer for visual cortex ROIs. The skull-stripped UNI volume was input to cat12, where SANLM denoising, bias correction and global and local intensity correction were performed before surface reconstruction. The local intensity corrected output of cat12 was supplied to recon-all with the hires pipeline and samseg segmentation. The cortical surfaces were then coregistered to the mean functional image using the OpenFmriAnalysis toolbox. First, the MT-weighted whole-brain EPI scan was coregistered and resliced with a rigid-body transformation to the mean functional using FSL FLIRT [64,65]. The whole-brain EPI was then used as the target for surface coregistration due to the improved grey/white matter contrast imparted by the MT-weighting. A rigid-body transformation between the UNI image and the whole-brain EPI was calculated using SPM12. This transformation was then applied to the coordinates of each vertex comprising the pial and white matter surfaces, resulting in cortical surfaces in the functional space of each participant. These surfaces were then used to define the cortical layers.

#### Definition of the cortical layers

The grey matter was divided into three equivolume layers using the level set method (described in detail in [66] and [67]) following the principle that the layers of the cortex maintain their volume ratio throughout the curves of the gyri and sulci [68,69]. The equivolume model transforms a desired volume fraction into a distance fraction, taking the local curvature of the pial and WM surfaces at each voxel into account [70]. We calculated two intermediate surfaces between the WM and pial boundaries, yielding three GM layers (deep, middle, and superficial). Based on these surfaces, we calculated four signed distance functions (SDF), containing for each functional voxel its distance to the boundaries between the five cortical compartments (WM, CSF, and the 3 GM layers). This set of SDFs (or “level set”) allowed the calculation of the distribution of each voxel’s volume over the five compartments [66]. For each cortical ROI (see below), voxels were assigned to one of the three GM layers only if >50% of its volume resided within that layer. This meant that any voxels with roughly equal proportion in each layer bin would not be selected.

#### Regions of interest

The medial temporal lobe was segmened using the automatic segmentation of hippocampal subfields (ASHS) [22] machine learning toolbox in conjunction with a database of manual 7T medial temporal lobe segmentations from a separate set of participants [23]. The cortical MTL regions were segmented with ASHS using an atlas of 3T MRI manual segmentations [28,29]. Hippocampal subfields from the 3T atlas were used in preliminary analyses before the 7T atlas became available and the results are presented in the supplemental information (Supplemental table 1). The hippocampal subfields and MTL cortices were defined on the anatomy of the T1- and T2-weighted images. The T1-weighted image was the denoised UNI volume and the T2-weighted image comprised the average of two high-resolution T2-weighted scans which were combined and denoised using the Structural Averaging Toolbox. All regions segmented with ASHS were coregistered to the mean functional image of each participant using FSL FLIRT. In addition, LOC was automatically defined on each participant’s T1-weighted image using FreeSurfer and coregistered to functional space using the transformation described above. All regions of interest were visually inspected for each participant and were collapsed over the left and right hemispheres as we had no hypotheses regarding hemispheric differences.

### fMRI analysis

#### fMRI preprocessing

All functional magnitude volumes were first denoised using the NORDIC denoising algorithm [71,72] implemented with the scripts available at https://github.com/SteenMoeller/NORDIC_Raw. Twelve slices were cropped from each EPI volume (six from the top and six from the bottom) due to low signal at the periphery of the slab excitation profile. The four blip-reversed (anterior-posterior phase encoding direction) volumes were removed from the time-series and used, together with the first four volumes (posterior-anterior phase encoding direction) to calculate the distortion-inducing B0 field with FSL Topup [62,73,74]. The topup field output was converted to a voxel displacement map using the Fieldmap SPM12 toolbox, which was used as input to SPM12 realign and unwarp for motion and distortion correction in one resampling step.

#### fMRI data quality

The temporal signal-to-noise ratio (tSNR) was computed by dividing the mean by the standard deviation of every voxel over the whole time-series of scans. We used the percentage of usable voxels, defined as voxels with tSNR > 5 divided by the number of voxels in the ROI, as a proxy measure of lost signal due to dropout. We then calculated the average tSNR for usable voxels. See Supplemental figure 1.

#### fMRI data modelling

The data from each run were analysed using a conventional GLM, with one regressor for each shape (separately for stimulus and omission trials in the prediction runs) and 18 head motion nuisance regressors. Shape-specific activity was calculated by subtracting the betas for shape B from shape A. Active voxels were chosen in visual cortex regions by selecting voxels with a t-value greater than 2 in a stimulus (shape B and shape B) vs baseline contrast. Voxels with less than tSNR of 5 were excluded from all regions based on the definition of “usable voxels” (see above). For layer-specific analyses, voxels were assigned to a layer only if more than 50% of their volume was Finally, for MTL regions, a cross-validation approach was used to select the most informative voxels based on the localiser run [7,8]. Voxels were first sorted by their shape-selective activity based on the t-value for a contrast of shape A vs shape B. Second, the multivoxel correlation was performed on different subsets of these voxels (between 10 and 100%, in 10% increments), within the blocks of the localiser run (trained on one run and tested on the other). For all iterations, the number of voxels that yielded the highest correlation value was selected. A multivoxel correlation analysis was performed with a Pearson’s correlation of shape-specfic activity patterns between the localiser run and prediction runs. Significance of the correlation for each region was determined by a one-sample t-test against zero for omission trials.

#### Informational connectivity

To investigate whether information is passed between the hippocampus and neocortex, we used an informational connectivity approach [8,31,32]. This approach infers connectivity from covariation in trial-by-trial multi-voxel pattern analysis (MVPA) measures between regions, and is therefore the MVPA analog of functional connectivity methods commonly applied to univariate data. Single-trial betas from the omission trials were correlated with the shape-specific activity of the localiser (see above) to gain a trial-by-trial measure of shape-specific information during prediction runs. Connectivity between hippocampal subfields and MTL cortical regions was performed by calculating the Pearson correlation between their timecourses while par-tialing out the the timecourses of all other subfields. As MTL regions are expected to covary due to spatial proximity, we also calculated scrambled pattern-similarity time courses by shuffling shape labels across 100 permutations to compare “baseline” and “real” connectivity. Hypothesis testing was performed with a two-way ANOVA with factors of layer (deep, middle, superficial) and connectivity (real, baseline).

## Acknowledgments

This work was supported by a Wellcome/Royal Society Sir Henry Dale Fellowship [218535/Z/19/Z] and a European Research Council (ERC) Starting Grant [948548] to P.K. The Wellcome Centre for Human Neuroimaging is supported by core funding from the Wellcome Trust [203147/Z/16/Z]. The authors thank Eleanor Maguire for valuable feedback on the results. We also thank Rosanna Olsen and Marshall Dalton for advice regarding hippocampal subfield segmentation.

## Supplemental information

### tSNR

**Supplemental figure 1:**
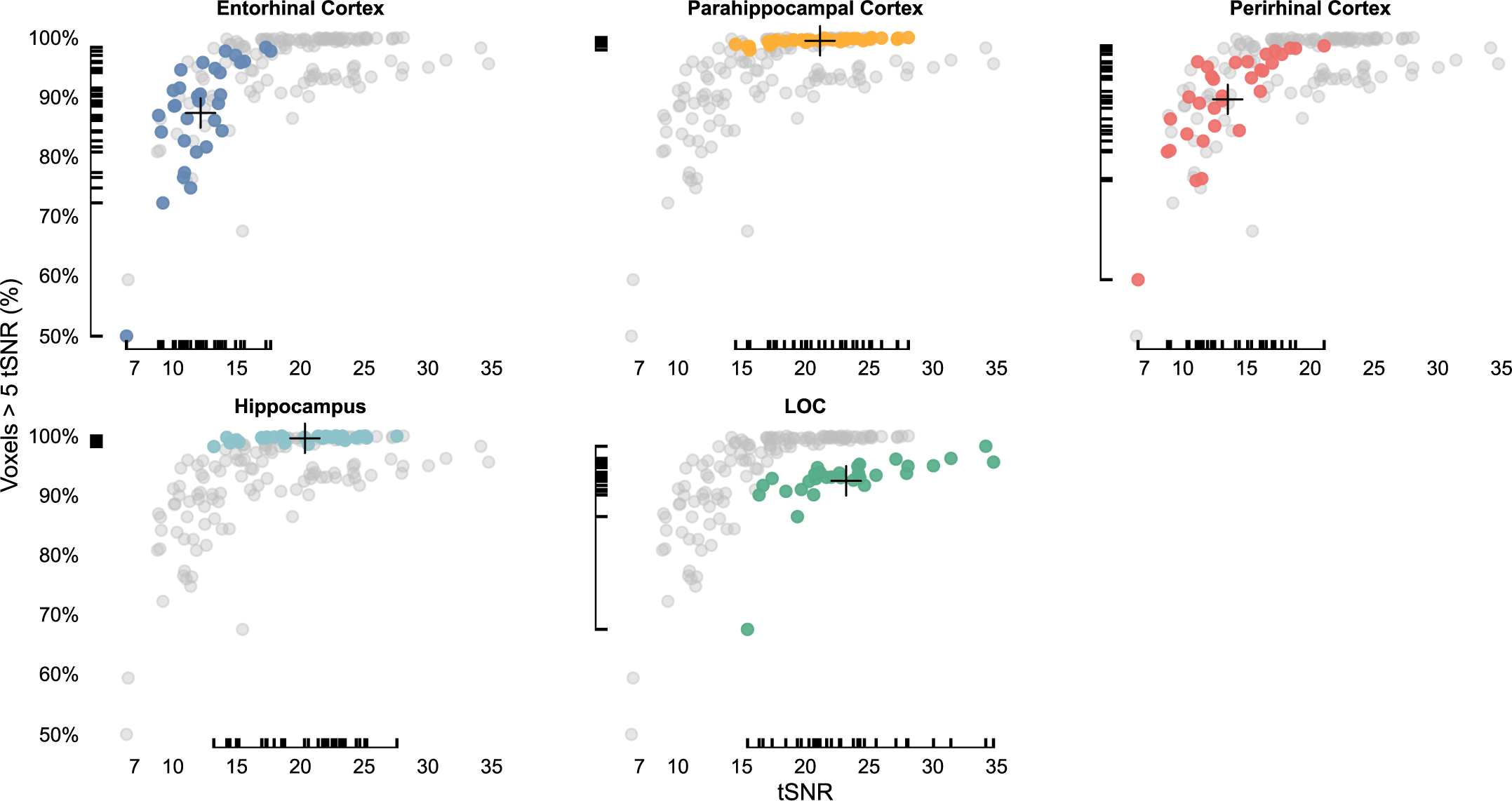
tSNR measures for each region of interest. For every subject, we measured the average tSNR and the percentage of usable voxels - voxels with tSNR > 5 - for each region. In each panel, the values for every subject in one particular ROI are highlighted in colour. To aid comparison, the values of all other ROIs are reproduced in gray for each panel. The mean ROI value of each axis are shown with a cross. Although the entorhinal and perirhinal cortices are considered challenging regions to scan [75], we developed a sub-millimetre 7T fMRI protocol in which they have reasonable, yet still lower, tSNR compared to our other regions of interest

### MTL Segmentation

**Supplemental figure 2:**
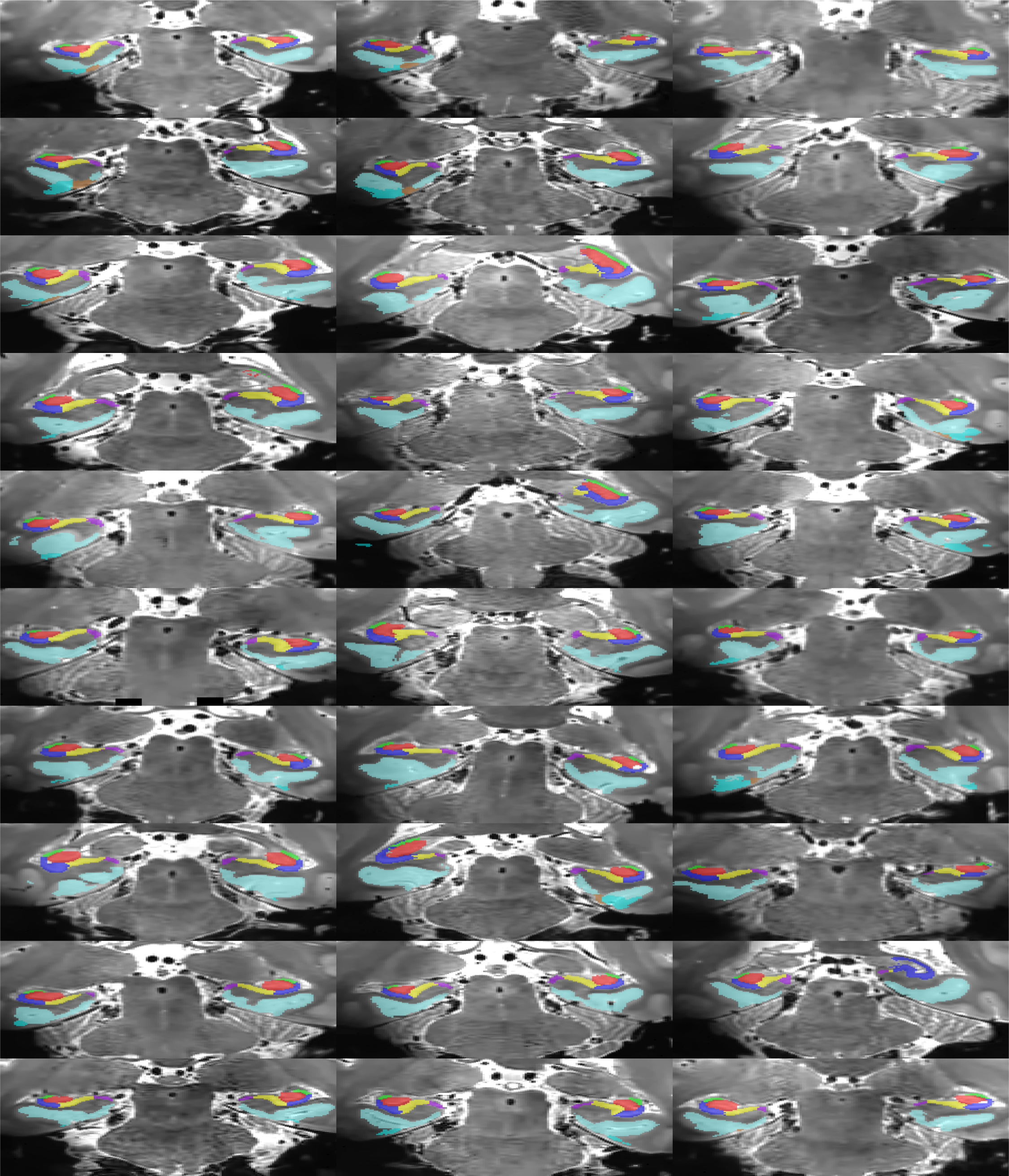
MTL segmentation overlaid on the high-resolution T2 image. Images show a coronal slice in the posterior hippocampus for each participant with the following ROIs visible: DG (red), CA23 (green), CA1 (blue), subiculum (yellow), pre/parasubiculum (purple), PHC (cyan).

**Supplemental figure 3:**
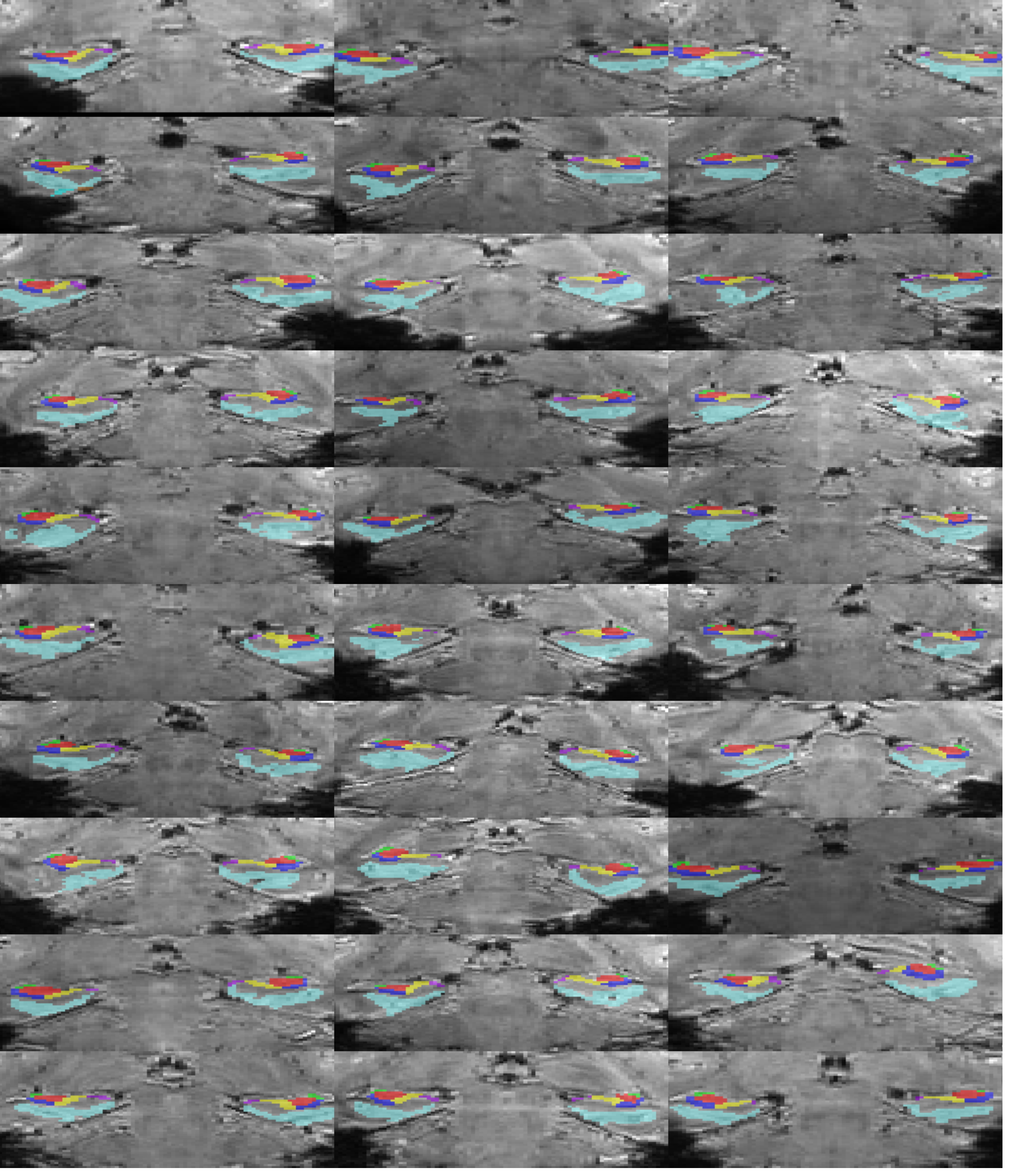
MTL segmentation overlaid on the mean functional image. Images show a coronal slice in the posterior hippocampus for each participant with the following ROIs visible: DG (red), CA23 (green), CA1 (blue), subiculum (yellow), pre/parasubiculum (purple), PHC (cyan).

### Entorhinal Cortex results

**Supplemental figure 4:**
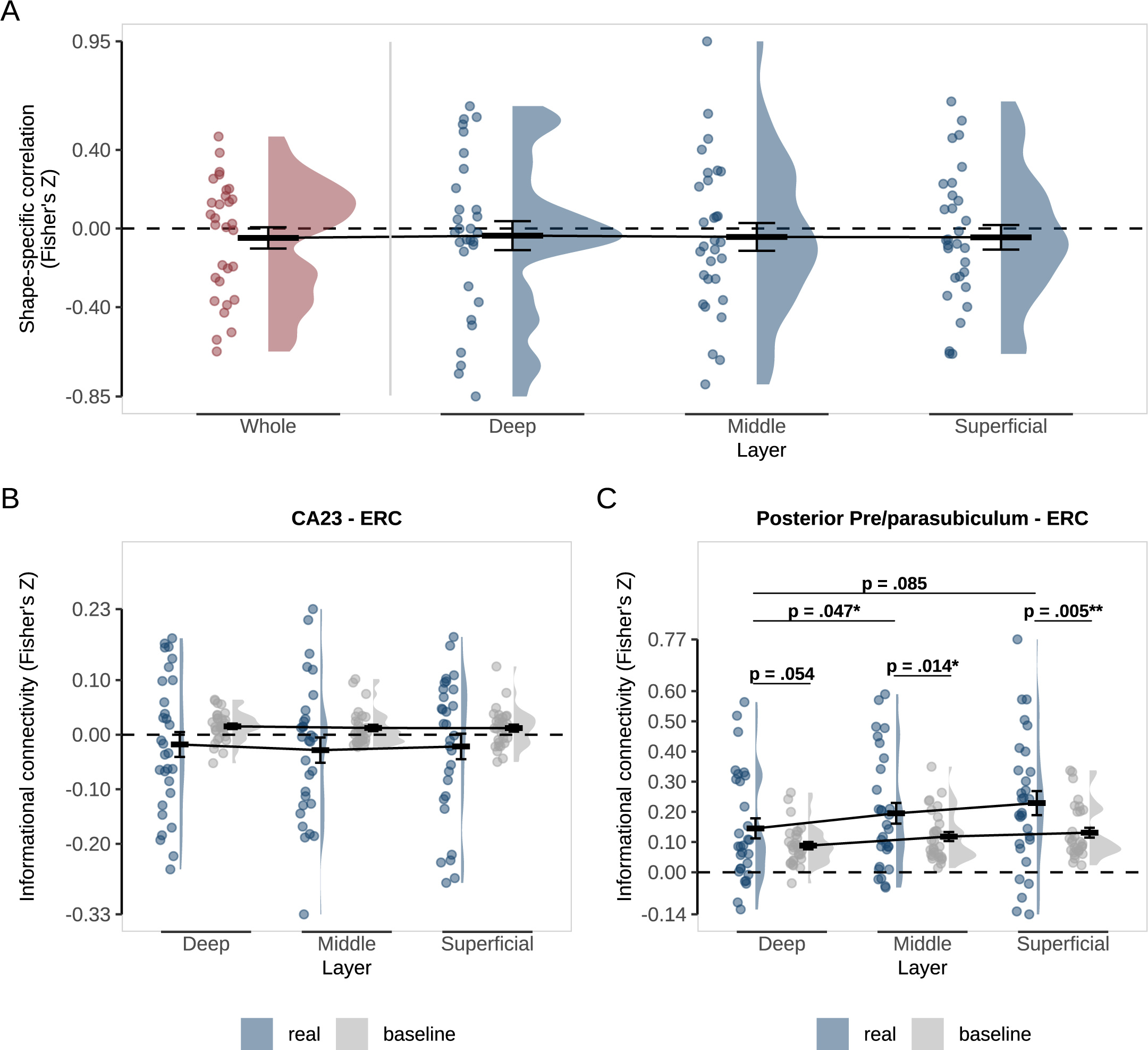
Representations and connectivity of ERC during omission trials. A) Pattern-similarity analysis in ERC as a whole (red) and specific to the deep, middle and superficial layers (blue). Pattern-similarity reflects the correlation between shape-specific (shape A - shape B) activity patterns in the localiser and omission trials. B) Informational connectivity of CA23 and C) pre/parasubiculum with ERC layers. Real connectivity (blue) is the observed correlation between regions on omission trials. Baseline connectivity (grey) was calculated by randomly shuffling the shape labels across 100 permutations. p values represent post-hoc paired t-tests investigating the differences between real and baseline and across layers. For all figures, crossbars and error bars represent the mean and within-subject standard error of the mean, respectively. Individual subject values are plotted in points alongside the probability density estimate.

### 3T ASHS atlas results

**Supplemental table 1:** Pattern similarity results for the 3T ASHS atlas [28,29]. In this study, we segmented the hippocampal subfields with the ASHS toolbox trained on atlases of manual segmentations. We initially used an atlas defined on 3T MRI data [28,29], but a 7T atlas became available during the analysis. The new atlas was generated with high-resolution T2-weighted scans collected on the same 7T scanner used in this study and provided a match in protocol and preprocessing [23]. Due to this match, we present the results from the 7T atlas in the main results. However, differences in segmentation between atlases were noticeable, such as the separation of the subiculum into pre/parasubiculum and subiculum proper in the 7T atlas, slight differences in the CA1-subiculum border and others. Therefore, we provide the results generated with the 3T atlas in the supplemental information for the interested reader. We wish to emphasise that we are making no claims about the quality of either atlas, especially as there is currently no universally agreed-upon protocol for subfield segmentation with MRI.

| Region | Z | t-value | p |
| --- | --- | --- | --- |
| Hippocampus | -0.06 | -1.58 | 0.126 |
| Anterior hippocampus | -0.02 | -0.55 | 0.585 |
| Posterior hippocampus | -0.10 | -2.23 | 0.034 |
| CA1 | -0.04 | -0.86 | 0.396 |
| Posterior CA1 | -0.08 | -1.63 | 0.114 |
| CA23 | -0.04 | -0.92 | 0.365 |
| Posterior CA23 | -0.04 | -0.92 | 0.365 |
| DGCA4 | -0.06 | -1.40 | 0.172 |
| Posterior DGCA4 | -0.04 | -0.79 | 0.434 |
| Subiculum | -0.09 | -2.12 | 0.042 |
| Posterior Subiculum | -0.19 | -3.63 | 0.001 |

